# Effect of glucose concentration in culture medium on the human preimplantation embryo methylome

**DOI:** 10.1101/2024.04.25.591072

**Authors:** Daniel Brison, Mollie McGrane, Sue Kimber

## Abstract

**Study question:** Does glucose concentration in culture medium have an impact on the DNA methylome of the early human embryo?

**Summary answer:** Glucose concentration is associated with changes in gene expression, global DNA methylation, methylation levels at CpG islands and at key histone modifications in human blastocysts.

**What is known already:** Preimplantation human embryos are highly sensitive to their local environment, and this may have long term implications for the health of the developing embryo, fetus and offspring. Glucose is a standard component of human embryo culture media, due to its importance as a nutrient. However, concentrations of glucose differ widely between different commercially available types. The present study was designed to determine whether changes in glucose concentration could influence global methylation and gene expression in the human preimplantation embryo.

**Study design, size, duration:** Human embryos were cultured in clinically relevant concentrations of glucose and global DNA methylation analysis was performed. The effect of glucose concentration on the embryo epigenome, specifically DNA methylation, was analysed.

**Participants/materials, setting, methods:** Human embryos surplus to treatment requirements were donated with informed consent from several ART centres. Embryos were cultured to the blastocyst stage in Vitrolife G-TL™ medium, either at 0.9 mM or 3.5 mM glucose, separated via immunosurgery into Inner Cell Mass (ICM) and trophectoderm (TE) samples, and compared for both DNA methylation and gene expression. This allowed us to evaluate the association between DNA methylation and previously importantly identified biological pathways.

**Main results and the role of chance:** The concentration of glucose in human embryo culture medium was associated with changes in gene expression and global DNA methylation in both ICM and TE, and methylation levels at CpG islands and key histone modifications. These results are significant because glucose is a major nutrient metabolised by human embryos in culture, and yet we know relatively little of its downstream effects on the genome and epigenome.

**Wider implications of the findings:** Commercially available embryo culture media with varying glucose levels have also been associated with altered fetal growth, birthweight and postnatal development of IVF offspring. Our findings may have important ramifications for potential clinical markers of embryo quality and pregnancy initiation, and improve understanding of the mechanisms underlying the impact of the early environment on the long term health of ART offspring.

**Study funding/competing interest(s):** This work was funded by the National Council for Science and Technology of Mexico (CONACyT), an NIHR pre-doctoral fellowship (PCAF) to MM, the NIHR Local Comprehensive Research Network and NIHR Manchester Clinical Research Facility, the University of Manchester and Manchester University NHS Foundation Trust. None of the authors has any conflict of interest to declare.

## Introduction

Following the first successful IVF treatment in 1978, the use of Assisted Reproductive Technology (ART) to treat infertility has risen significantly, with over 9 million children now born as a result (ESHRE, 2021). The development of ART has increased the understanding of human preimplantation development, including how the embryo culture environment can have an impact (Gardner and Kelley, 2017). During development, the initial preferred energy source for human preimplantation embryos is pyruvate, as the embryo relies on oxidative metabolism to obtain ATP. At the morula stage, there is a switch to glucose uptake due to increased hexokinase activity, leading to an increase in oxygen consumption (Brison and Leese, 1991; Leese, 2012). Substantial alterations in these pathways can compromise embryo metabolism, potentially affecting embryo development and viability. Within a clinical setting, the general composition of embryo culture media is modelled on these early embryo energy requirements (Cairo Consensus Group, 2020). However it is known that the relative concentrations of nutrients present in ART media can directly affect embryo metabolism (Gardner et al., 2013). Embryo culture media manufacturers are not legally required to disclose the concentrations of constituents of their media, so the precise compositions of their products are still unknown. However, research has identified wide variations in essential ingredients such as glucose, pyruvate, lactate and amino acids, between different commercially available media (Morbeck et al., 2017). The true clinical implications of these differences are undetermined, highlighting the importance of extended research in this area.

According to the developmental origins of health and disease (DOHaD) hypothesis, unsuitable environmental exposure in utero has the potential to predispose the offspring to certain diseases in adult life (Feuer and Rinaudo, 2012). The mechanism of this is thought to be driven by changes to the epigenetic profile of the developing embryo (Mandy and Nyirenda, 2018). Epigenetic modifications have the ability to regulate gene expression without altering the DNA sequence (Fleming et al., 2015). Preimplantation embryos experience profound reprogramming of epigenetic information inherited from the gametes; this is an essential process during early embryo development (Ivanova *et al*., 2020; Wilkinson et al 2023). DNA methylation plays a large role in this epigenetic event, whereby a global wave of DNA methylation occurs to ultimately secure totipotency and pluripotency in the embryo (Arand et el., 2021). ART interventions overlap with this crucial period, meaning that exposure to the artificial *in-vitro* environment could potentially affect these pathways. Specifically, DNA methylation is known to be a malleable genetic mark, as it can be susceptible to environmental influence. Studies have demonstrated that certain ART interventions can affect the levels of DNA methylation in mouse embryos (Wright et al., 2011) and in human placentas (Ghosh et al., 2017). The long-term impacts of this are yet to be fully understood, but may impact gene expression regulation, establishing imprinting patterns and maintaining genome stability (Li et al., 2013; Wilkinson et al 2023; Ho et al., 2016). Furthermore, it has been recorded that offspring conceived via ART display differences in DNA methylation levels, when compared to naturally conceived offspring (Håberg et al., 2022). Therefore, it is important to establish which aspect of ART may be impacting embryo epigenetics. These changes could in part be linked to the inherent infertility of one or both parents (Pinborg et al., 2013), but artificial culture conditions are likely to play a role. Whilst implantation and live birth rates do not seem to be strongly associated with medium type, several metrics of offspring growth are; the speed of post-implantation embryo development (Van Duijn et al., 2020), the initial birth weight of offspring (Dumoulin et al., 2010, Eskild et al., 2013) and the weight/body composition of children at age 9 (Zandstra et al., 2018) can all differ when comparing offspring born via different ART media types. It is therefore essential to gain more knowledge around which constituents within the culture media could be contributing to these changes, and particularly whether they are contributing through DNA methylation.

Glucose is a standard component of human embryo culture media due to its importance as a nutrient. However, concentrations of glucose differ widely between different commercially available types (Morbeck et al., 2014; 2017). The present study was designed to determine whether changes in glucose concentration could influence global methylation and gene expression in the human preimplantation embryo. We also aimed to analyse whether these changes were associated with altered signalling pathways that could affect embryo development.

To address this aim, human embryos were cultured in clinically relevant concentrations of glucose and global DNA methylation analysis was performed. The effect of glucose concentration on the embryo epigenome, specifically DNA methylation, was analysed. Embryos cultured in Vitrolife G-TL™ medium, either at 0.9 mM or 3.5 mM glucose, were compared for both DNA methylation and gene expression. This allowed us to evaluate the association between DNA methylation and previously importantly identified biological pathways.

## Materials and Methods

### Study design

This study utilised fresh and frozen human embryos, donated by couples undergoing ART treatment at St Mary’s Hospital in Manchester. Ethical approval was obtained by the Central Manchester Research Ethics Committee, under licence R0026 from the Human Fertilisation and Embryology Authority. All embryos were graded by a qualified Clinical Embryologist following the Alpha/ESHRE 2011 guidelines (Alpha Scientists in Reproductive Medicine and ESHRE Special Interest Group of Embryology, 2011).

Slow frozen embryos were thawed using the THAWKIT™ CLEAVE (Vitrolife, Sweden) following the manufacturer’s instructions and clinical standard operating procedures. Post thawing/warming, embryos were placed in 0.5 ml of pre-equilibrated GTL medium, overlaid with OVOIL™ (Vitrolife, Sweden) and incubated to allow for recovery and blastocyst formation.

To obtain clinically relevant results, embryos were cultured in the same conditions used in the Assisted Reproductive Technology (ART) clinic; 37°C, 6% CO_2_ and 5% O_2_. For the low glucose condition, we utilised G-TL™ medium (0.9mM glucose). For the high glucose condition, G-TL™ medium was supplemented with additional glucose to reach the concentration of 3.5 mM. The selection of glucose concentrations was based on data from the Morbeck group, regarding composition of commercial embryo culture media (Morbeck *et al*., 2014; 2017). In this case, 0.9 mM glucose was an average concentration and 3.5 mM was one of the highest concentrations used for clinical human embryo culture.

All reported significant results were calculated as p<0.05 after Benjamini and Hochberg correction, with a minimum difference of 10.0.

### Embryo preparation for analysis

Human embryos were collected and divided into two groups, comprising of 5 embryos in each group. Embryos assigned to the low glucose group were cultured for 6 days in standard G-TL™ medium (0.9mM glucose). Embryos assigned to the high glucose group were also cultured for 6 days but using standard G-TL™ medium (0.9mM glucose) from day 1 to day 3; on the morning of day 3, a media change was performed into the modified G-TL™ medium (3.5Mm glucose). On day 6, blastocysts were graded and dissected by immunosurgery to isolate the ICM and TE. Samples were isolated separately, either for DNA methylation analysis or gene expression evaluation.

### Evaluation of gene expression

Embryos were cultured from the pronuclear (PN) stage (day 1) until the blastocyst stage (day 6), where they were then processed through the PolyA-PCR protocol (Brady and Iscove 1993). We analysed the effect of glucose concentration on markers of apoptosis, pluripotency, metabolism, mitochondrial activity, hypoxia, oxidative stress and methylation, among others. Gene expression analysis was undertaken using the comparative CT method for semi-quantitative real-time PCR data (2^-ΔΔ CT^ method) using β-actin as the reference gene. We examined the expression of 48 genes including markers for: cell survival (BAX, BCL2, BCL2L1, CBL, DAP3, TP53); cell growth and differentiation (CDX2, EGFR, EOMES, EP300, FGF4, ZSCAN4); pluripotency (NANOG, OCT4, SOX2); transcription factors (EIF1AX, EIF2S2, TRIM28); DNA methylation (DNMT1, DNMT3A, DNMT3B, TET1, TET2, TET3, MAT2A); mitochondrial activity (ATP5F1, COX11, SDHB, TFAM TIMM23, TRMT10C); metabolism (SLC2A1, SLC2A3, SLC7A3, SLC16A1, GAPDH, ASNS, MTOR, PPARA, PPARG).

### Library preparation (PBAT protocol; Smallwood et al, 2014)

In order to complete library amplification, samples went through several processes: DNA extraction, bisulphite conversion, desulfunation, first and second strand synthesis as well as several purifications. DNA extraction was completed using an adapted EZ-DNA Methylation kit (D5001, Zymo Research). Total sample volume from immune-surgery process (∼5 µL) was mixed in a PCR tube with 19 µL master mix and incubated for 60 min at 37°C. Bisulphite conversion was completed by adding 125 µL of conversion reagent to each sample and incubating at 98°C for 8 min, followed by 64°C for 3.5h.

Bisulphite converted samples were desulphonated and purified using Zymo-spin IC columns, previously treated with 300 µL of M-Binding buffer. Converted samples were transferred to the column, followed by extra 100 µL of M-Binding buffer; this was used to wash and collect residuals from the previous reaction tube. Columns were centrifuged for 30 sec at 13000 rpm. 100 µL M-Wash buffer was added to the column, followed by centrifugation 30 sec/13000 rpm. 100 µL M-Desulphonation buffer was added and incubated for 20min at room temperature (RT). The column was then centrifuged 30 sec/13000 rpm and flow through was discarded. The column was washed twice with 200 µL of M-Wash buffer and then transferred into a new clean 1.5 mL tube. 40 µL of Nuclease free water was added to the column, incubated at RT for 5 min and centrifuged at 13000 rpm/ 1 min. The eluted flow-through was transferred to clean PCR tubes and mixed with first strand master mix, composed of 5 µL of NEB Buffer 2 (10X), 2 µL dNTP mix (10 µM) and 2 µL of 10 µM 6N-F primer (5’-CTA CAC GAC GCT CTT CCG ATC TNN NNN N-3’). Samples were incubated at 65°C for 2 min and immediately cooled down on ice; 1 µL of Klenow-exo enzyme (50 U) was then added. Tubes were incubated for 5 min at 4°C, 37°C for 90 min (rampage 1°C/ 15 sec) followed by 4°C hold. First strand synthesis was purified using AMPure XP beads (Beckman Coulter, US) at 1:0.8 ratio. 40 µL AMPure XP beads were added, mixed and incubated for 10 min at RT. After incubation, the reaction tubes were place on a magnet and samples were washed twice with 100 µL 80% Ethanol. Tubes were left open for 7-10 min until the beads were dry and the 48 µL of second strand master mix was added, to proceed with second strand synthesis. Second strand synthesis mix was incubated at 98°C for 1 min and immediately cooled to 4°C on ice. Two microliters of Klenow exo- (50 U) was added to each sample; this was mixed, briefly spun down and immediately transferred to the thermocycler. Samples were incubated as follows: 5 min at 4°C, then raised to 37°C with 1°C per 15 sec (33x ramp steps), followed by for 90 min at 37°C. Samples were vortexed every 20-30 min to re-suspend the beads, and were kept at 4°C after finishing the incubation. Second strand synthesis was purified by adding 50 µL of nuclease free wate and 80 µL ofAMPure XP buffer. Samples were mixed thoroughly and incubated for 10 min at RT. After incubation, samples were placed on a magnet and washed twice with 180 µL of 80% ethanol. Samples were then incubated at RT for 10 min until the beads were dry and they were eluted with 49 µL of Library amplification master mix. 1 µL of specific iTAG was added to each sample and they were then transferred to a thermocycler to proceed with the library amplification. Amplified libraries were purified by adding 35 µL of AMPure XP buffer to each product. They were well mixed by pipetting up and down, and then incubated for 10 min at RT. After incubation, libraries were placed on a magnet and washed twice with 100 µL of 80% ethanol. Libraries were then incubated at RT for 7-10 min until the beads were dry and then eluted in 15 µL of Nuclease-free water, where they were incubated for 2 min at RT. Tubes were then placed on a magnet; carefully, 13-14 µL of supernatant was transferred to clean tubes, without taking any beads. Purified libraries were stored at -20°C for further analysis.

### Global Methylation Analysis

The experimental procedure was performed on ICM and TE samples, but samples were paired and treated as single embryos during data analysis. Once libraries were synthesised and purified, they were analysed using capillary electrophoresis to determine the mean fragment sizes. Once the size was determined, all samples were normalised and quantified immediately before the sequencing analysis.

We aimed to identify whether any genes (and potential associated biological processes) were altered during preimplantation development, by the concentration of glucose in the medium. To address this, we first identified genes that were overlapped by probes defining differentially methylated regions. To identify candidate loci, we searched for regions that were differentially methylated in high glucose samples compared to low glucose. Data analysis and visualisation were performed using the SeqMonk software (RRID:SCR_001913), which relies on R and Java scripts to create plots. Cytosine and Guanine (CpG) methylation were calculated as the average of methylation for each CpG position. Comparisons to obtain hypermethylated and hypomethylated regions were set as those with a minimum difference of 10% in the methylation level. Samples were analysed using the EdgeR test, with all significant results recorded as p<0.05, alongside a significant false discovery rate (FDR). Differentially methylated genes were evaluated via ORA and WebGestalt.org.

Global methylation analysis was assessed by comparing high glucose and low glucose paired combined samples (ICM+TE). For the first global methylation analysis, methylated points (MPs) were mapped according to the CG_occurrences_GRCh38.txt.gz database. Subsequently, methylated regions (MRs) were defined through the genome by spanning probes over 100 CpGs, which resulted in 288,333 probes. Unbiased measures of methylation were performed over a fixed window size of 100 methylated points (CpGs) as this was found to give at least 100 reads per methylated region.

### H3K4me3 + H3K27me3 Sites Analysis

Potentially active promoters were mapped using published data sets (accession number GSE124718) for ChIP H3K4me3 on ICM. Called picks on ChIP-seq analyses were used to guide selections of regions within the blastocysts (ICM + TE) genome, for further evaluation. From these mapped regions, differentially methylated genes between the low and high glucose groups were identified. Groups were compared using EdgeR test, with the filtering criterion of adjusted p value (p <0.05) and difference in methylation greater than 10% (Δβ ≥10percentage).

### Differential Methylation of CpG Islands

Bisulfite pipeline (Bismark) was used for quantification of methylation levels and EdgeR was used for the statistical analysis (p<0.05) plus Benjamimi and Hochberg correction, with a minimum difference of 20.0%.

### Imprinted Genes

We investigated the effect of glucose concentration on previously reported imprinted human genes in our samples. Imprinted genes were mapped using SeqMonk software (Homo sapiens genome GRCh38) according to detailed location from the imprinted library from the University of Otago (https://www.otago.ac.nz). Every imprinted gene was counted as a whole potential methylated region, independently of the methylated points (CG occurrences) contained per region. Bisulfite pipeline (Bismark) was used for quantification of methylation levels and EdgeR was used for the statistical analysis.

## Results

### Effect of glucose concentration on blastocyst gene expression

Key pluripotency-associated transcription factors, OCT4 and SOX2, showed differential expression between the high and low glucose groups (Fig. 2). Both markers were significantly downregulated in low glucose, compared to the high glucose condition.

**Figure 1:**
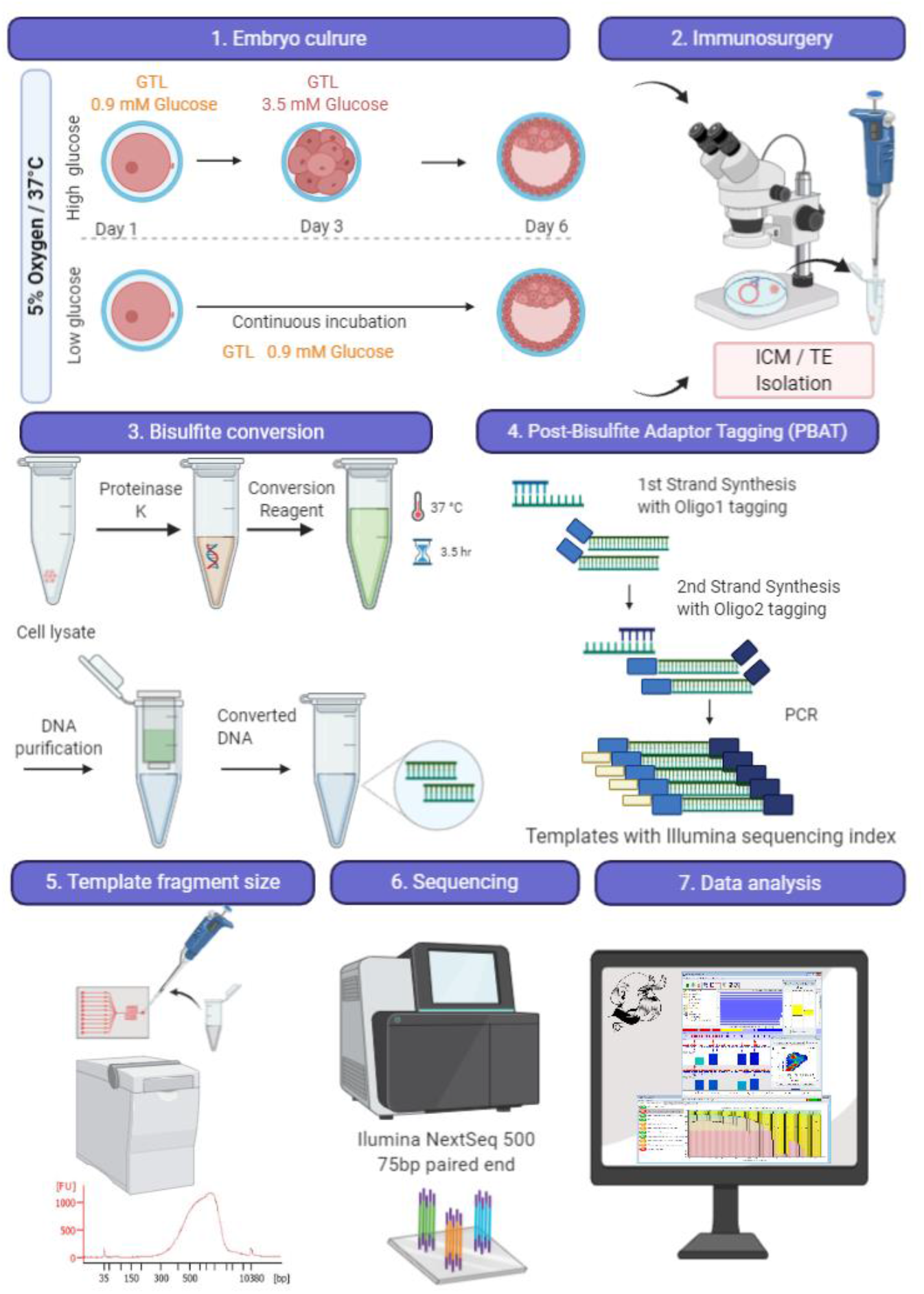
DNA Methylation analysis of human preimplantation embryos. 1) All embryos were cultured in G-TL™ with 0.9mM glucose from day 1 to day 3 morning. Embryos in the high glucose group were then switched to 3.5mM glucose in G-TL™ from day 3 to day 6. 2) ICM and TE was isolated from day 6 blastocysts. 3) DNA was isolated and treated with bisulphite 4) BS-seq libraries were prepared through the post-bisulfite adapter tagging method. 5) Library quantification was assessed using high-sensitivity DNA chips on the Agilent Bio-analyser. 6) Pools of 10 libraries were prepared for 75bp paired-end sequencing on a NextSeq500 and run in duplicate. 7) Unbiased analysis was performed in SeqMonk.

**Figure 2:**
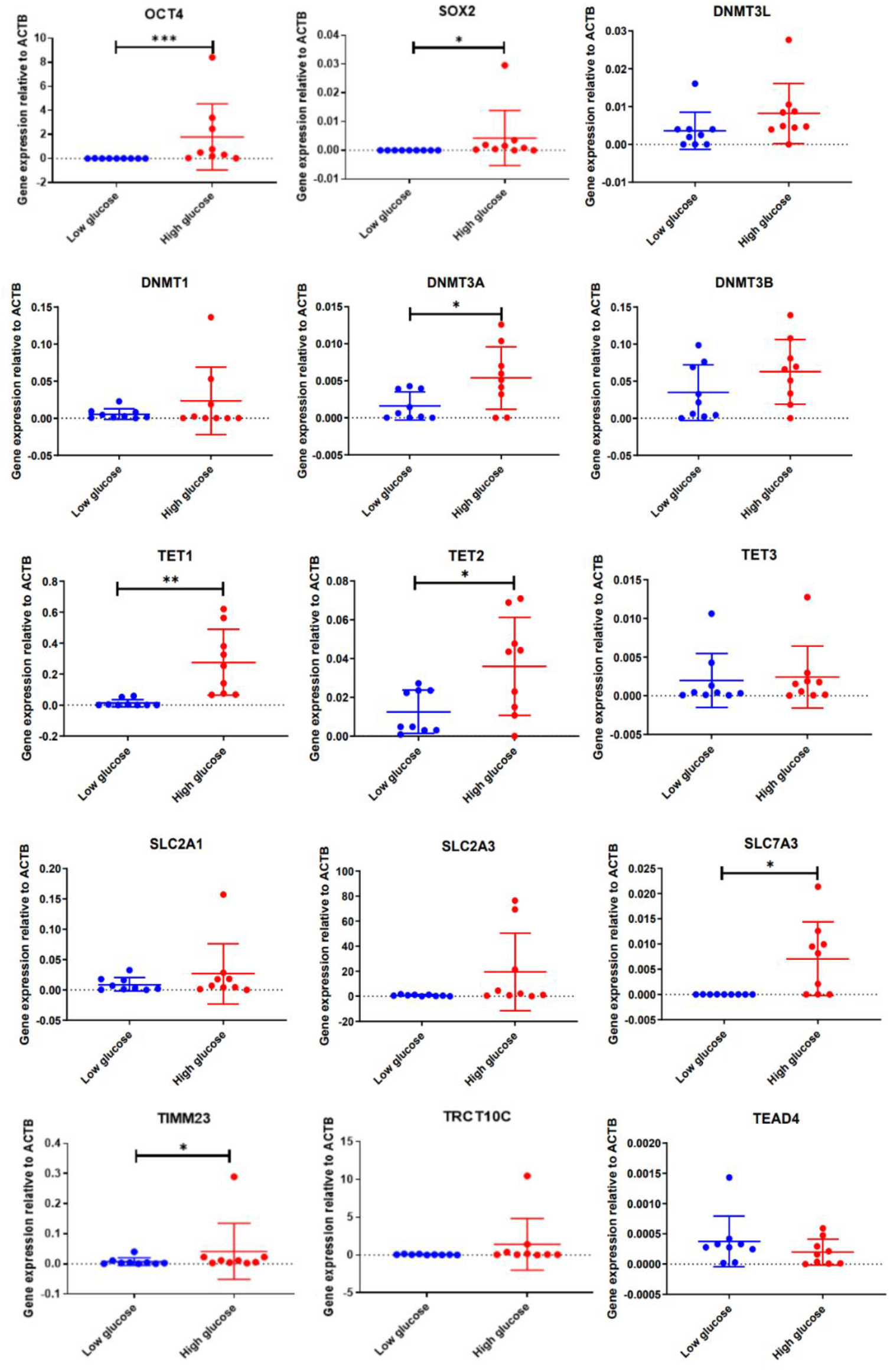
The effect of glucose concentration on blastocyst gene expression. Expression of pluripotency-associated genes (OCT4, SOX2), DNA methyl-transferase genes (DNMT1, DNMT3A, DNMT3B, DNMT3L), DNA demethylases (TET1, TET2, TET3), glucose transporters (SLC2A1, SLC2A3), amino acid transporter (SLC7A3), mitochondrial importer (TIMM23), Mitochondrial RNase P (TRCT10C) and the Hippo pathway component (TEAD4) expression in human embryos cultured for 6 days in either 0.9 mM glucose (low glucose) or 3.5 mM glucose (high glucose). The data plots show the mean expression +/-standard deviation (SD) of three technical replicates per sample. Significance (p<0.05).

Mitochondrial RNase P (TRCT10C) expression was not significantly different between the two groups, whereas the mitochondrial importer (TIMM23) showed significantly higher expression in the high glucose group (Fig. 2).

DNA methyltransferase activity was variable between the groups; the maintenance methyltransferase (DNMT1) and the de novo methyltransferases (DNMT3B and DNMT3L) did not show any significant differences between the glucose conditions. However, the de novo methyltransferase DNMT3A showed an upregulated expression in the high glucose condition. Demethylases TET1 and TET2 also showed increased expression in high glucose, whereas TET3 did not show any significant difference. Glucose transporters SLC2A1 and SLC2A3 were not differentially expressed but amino acid transporter SLC7A3 was significantly upregulated in high glucose. Moreover, the transcription factors ZSCAN4, EIF2S2 and EOMES (not shown), and the Hippo pathway component, TEAD4, did not show significant changes between the high and low glucose groups.

### Global methylation in blastocysts in low glucose vs high glucose

Having noted differences in gene expression (notably in DNMT3A TET1 and 2) in blastocysts, associated with glucose level, we analysed global DNA methylation levels in blastocysts. Methylated regions (MRs) were defined through all of the genome by spanning probes over 100 CpGs.

The average levels of methylation (0-100%) and percentage of abundance in the two groups are shown (Fig. 3). Interestingly, the high glucose group had a greater number of methylated genes relative to the low glucose group, with 23% of the genome presenting from 50-75% methylation.

**Figure 3:**
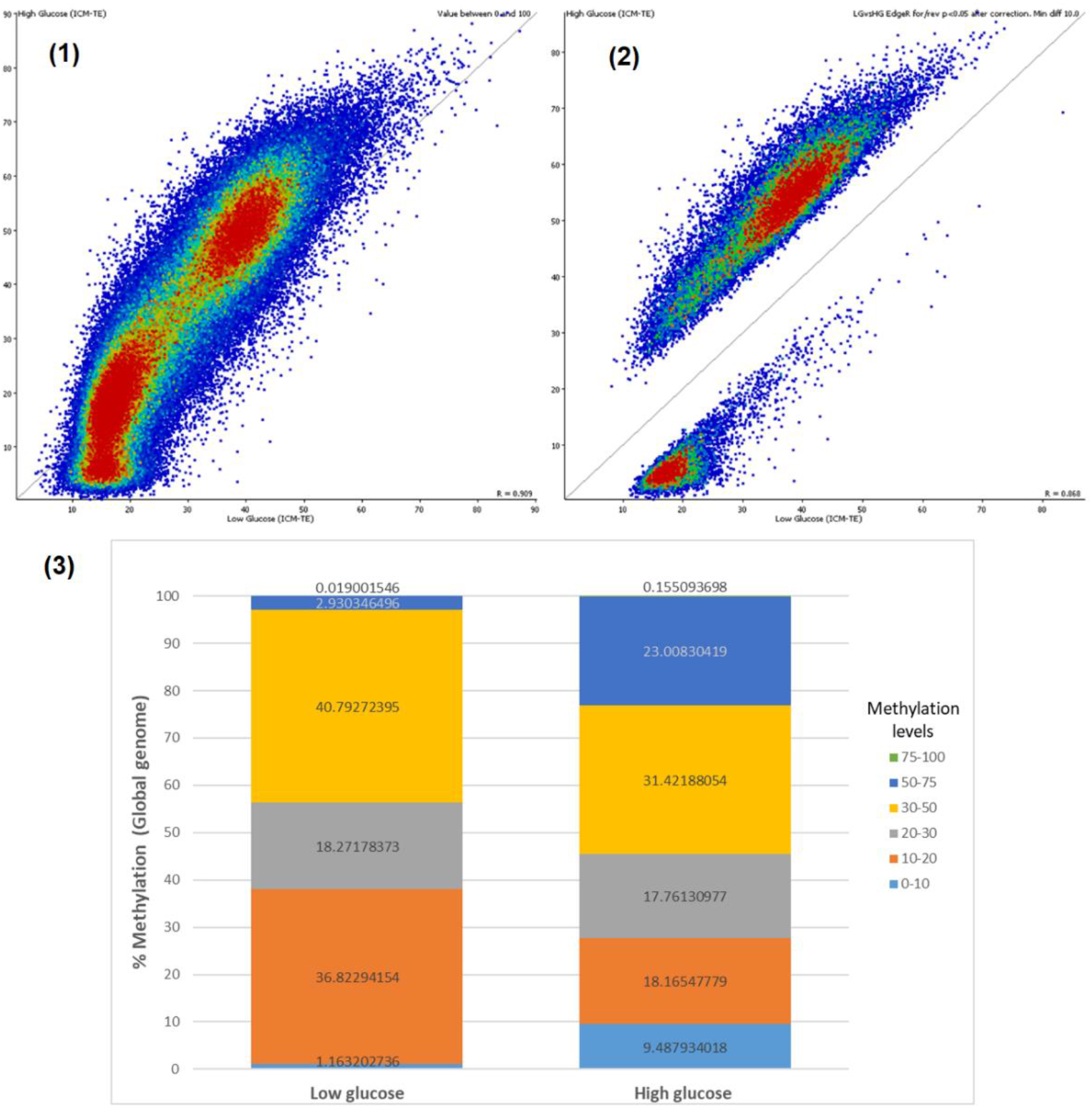
Correlation between low glucose ICM+TE and high glucose ICM+TE. **(1)&(2):** Scatter plots showing the correlation between data groups of merged samples: low glucose ICM+TE (x-axis) and high glucose ICM+TE (y-axis). Each dot represents a methylated region (MR) spanning 100 methylated points (CpGs). **(1):** Overall correlation of methylation levels for 285517 MRs in merged ICM+TE samples. The plots show a common scale for methylation levels (0-90%) in both data sets and a Pearson correlation value R=0.909. **(2)** Correlation of 57737 DMRs resulted after EdgeR statistical test (p=<0.05, Δβ≥10%). The plots show a common scale for methylation levels (0-80%) in both data sets and a Pearson correlation value R=0.868. (3): Distribution of methylation levels of human blastocysts between the two glucose conditions; the y-axis is indicating the percentage of abundance from 0-100. Bars are representing the total (288,333 = 100%) of methylated regions per group. Each colour indicates a range of methylation levels and the numbers within the bars/colour are indicated the abundance of methylated regions.

Groups were compared using EdgeR test with the filtering criterion of adjusted p value (p <0.05) and difference in methylation greater than 10% (Δβ ≥10%). This resulted in 57,737 probes identified as being differentially methylated; of which, 18.1% were hypomethylated and 81.9% hypermethylated in high glucose (Fig. 3, Fig. 4). The mean methylation of gene bodies from both groups was also compared using a quantitation trend plot, showing the relative methylation levels of gene bodies (±5000bp) in low glucose and high glucose groups (Fig. 4). Here, we confirmed that the high glucose group showed greater mean methylation levels, relative to the low glucose samples.

**Figure 4:**
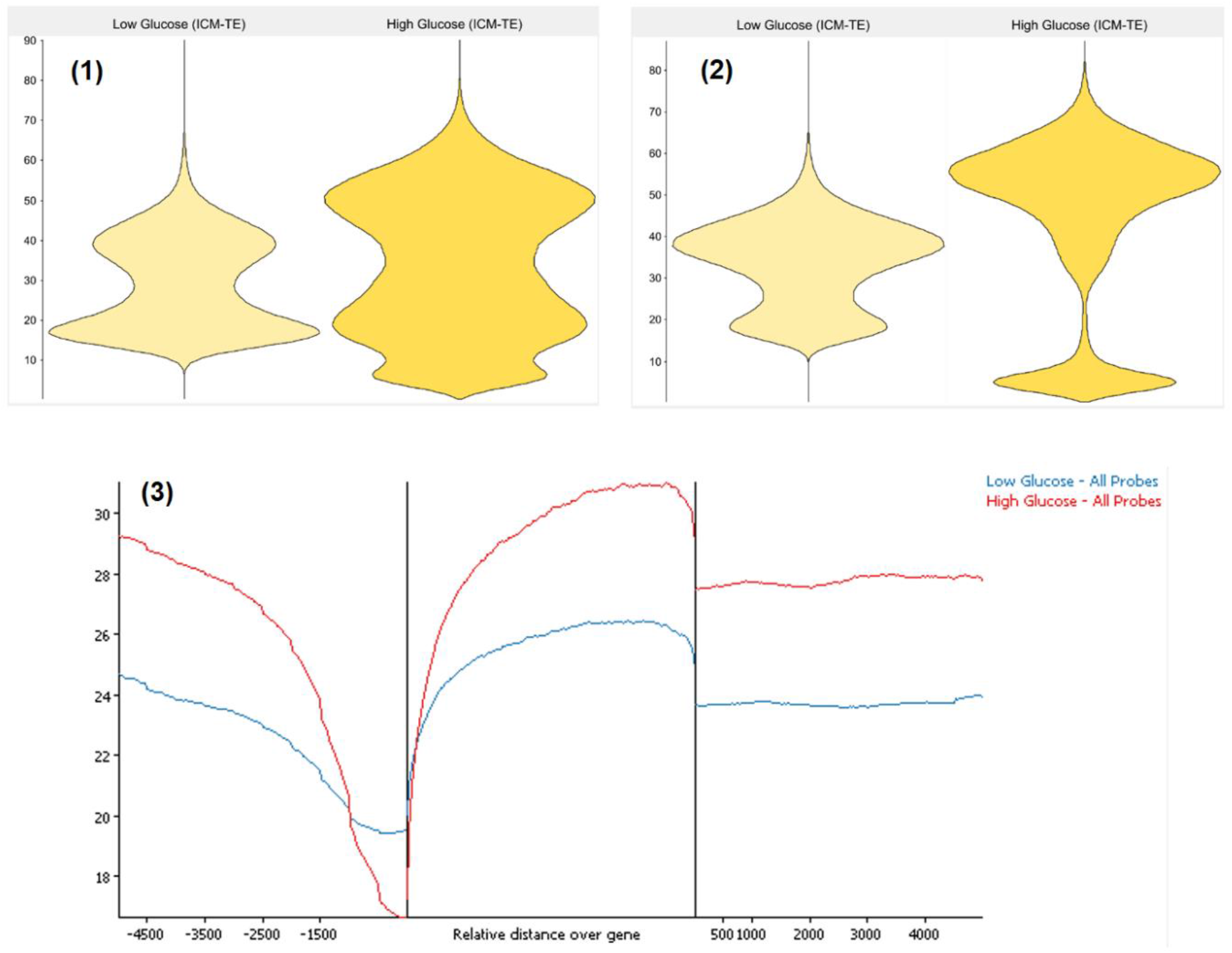
**(1):** Bean plot showing the mean distribution of the global methylation levels (0-90%) in the low glucose (n=5) and high glucose (n=5) groups, using patient-paired samples (5* ICM-TE) Only values above zero have been shown. **(2):** Bean plots showing the mean distribution of DMRs in the low glucose and high glucose groups. DMRs were identified after applying the adjusted EdgeR test (p <0.05, Δβ ≥10%): 18.1% were hypomethylated and 81.9% hypermethylated, when comparing the two groups. **(3):** Quantitation trend plot, showing the relative methylation levels of gene bodies ±5000bp in low glucose (blue) and high glucose (red) groups. This representative plot shows the mean distribution of a gene body, including its upstream and downstream regions (±5000 bp). Overall, low glucose samples are hypomethylated in both the upstream region and downstream region of gene bodies, in comparison with the high glucose samples. Interestingly, there is a small region about -500 to -1000 bp where the high glucose samples become hypomethylated.

We next defined 8654 genes as being differentially methylated (DM), when overlapped by at least one probe containing 100 CpGs (Fig. 3). Then, a subset of genes whereby methylation levels decreased in high glucose or increased in low glucose (and vice versa) were identified. Results from ORA, from 8654 differentially methylated genes, showed enriched gene sets related to the cell cycle, embryo development, regulation of gene expression and intracellular transport. Disease associations generated from the OMIM database included leukaemia, breast cancer, diabetes, mitochondrial deficiency and Leigh syndrome.

### Methylation in ICM and TE in high and low glucose

Low and high glucose samples clustered apart independently of ICM/TE cell type, suggesting that glucose may be affecting methylation in both (Fig. 5). The ICM was also compared to the TE for each glucose concentration.

**Figure 5:**
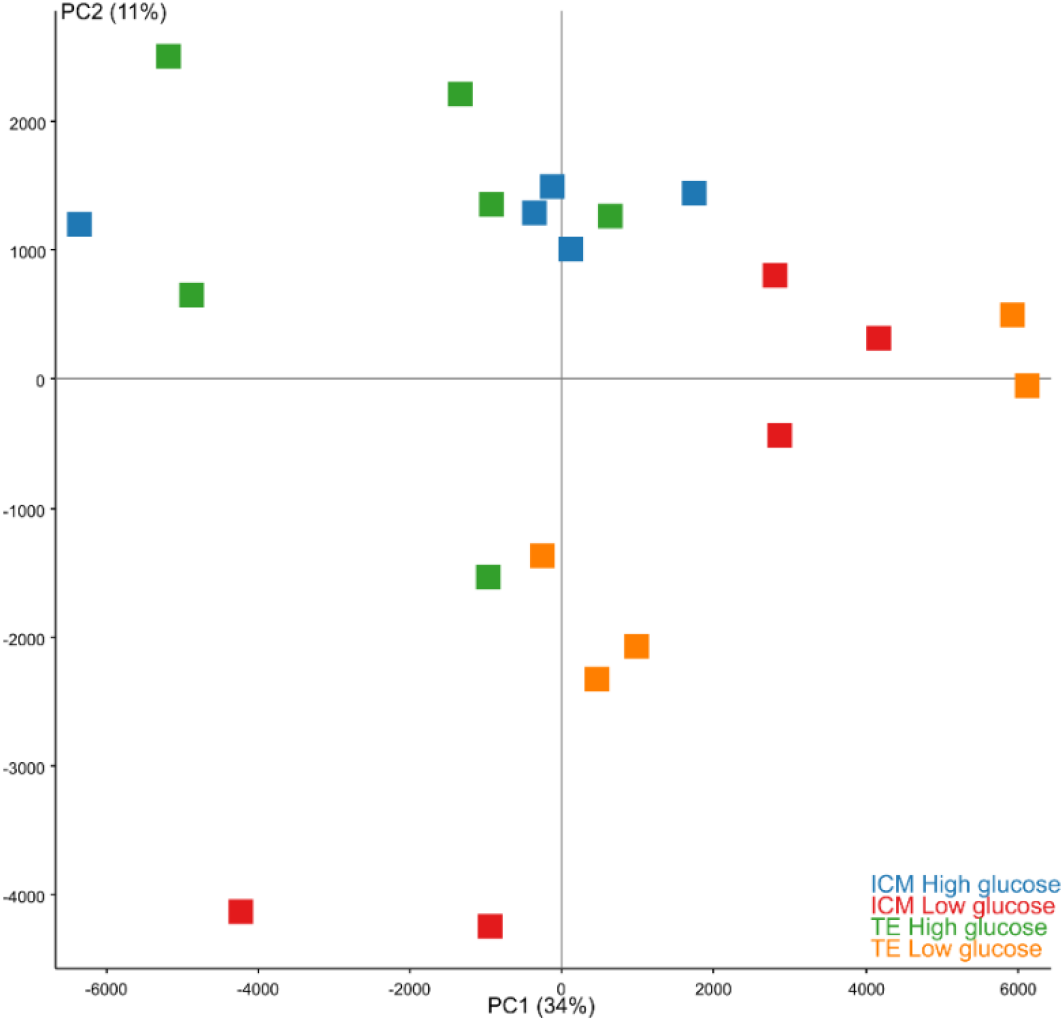
PCA plot of DNA methylation profiles of high and low glucose samples (isolated ICM and TE). Each coloured dot represents an individual DNA sample from high glucose ICM (blue), low glucose ICM (red), high glucose TE (green), or low glucose TE (orange) groups.

### DNA methylation in ICM in low glucose vs high glucose

A significant difference was found when comparing the ICM in high glucose vs low glucose. There were 5023 differentially methylated regions (DMRs), which were overlapping 2755 genes. Methylated regions from the low glucose group were plotted against methylated regions from the high glucose group, with a correlation of 0.872.

Of the 2755 differentially methylated genes that were identified, the most enriched gene sets were related to regulation of biosynthetic processes within the cell, regulation of gene expression and neurogenesis. Regarding disease terms, using the OMIM database, cancer was the most common output, followed by heart malformation, mitochondrial deficiency and obesity. From the GLAD4U database, the most enriched diseases were related to cell adhesion processes.

151 hypermethylated genes within the ICM in the high glucose group were identified. Although the gene ontology results were significant, false discovery rate (FDR) was not (data not shown).

In the ICM, 2604 significantly hypermethylated genes in the low glucose group were identified, with both the gene ontology results and FDR significant. Here, the most enriched gene sets were related to regulation of biosynthetic processes, gene expression and neurogenesis. Regarding diseases related to these genes found using the OMIM database, cancer was again the most common output, followed by heart malformation, mitochondrial deficiency and obesity. When using information from the GLAD4U database, the most enriched diseases were related to cell adhesion, cell stress and cancer.

### Differences in DNA methylation between TE in low glucose vs high glucose

A significant difference was determined between TE samples in high glucose compared to low glucose. This resulted in 25234 DMRs, which were overlapping 5079 differentially methylated genes (DMGs). DMRs were plotted for correlation analysis and to visualise abundance and mean methylation levels (Fig. 6).

**Figure 6.**
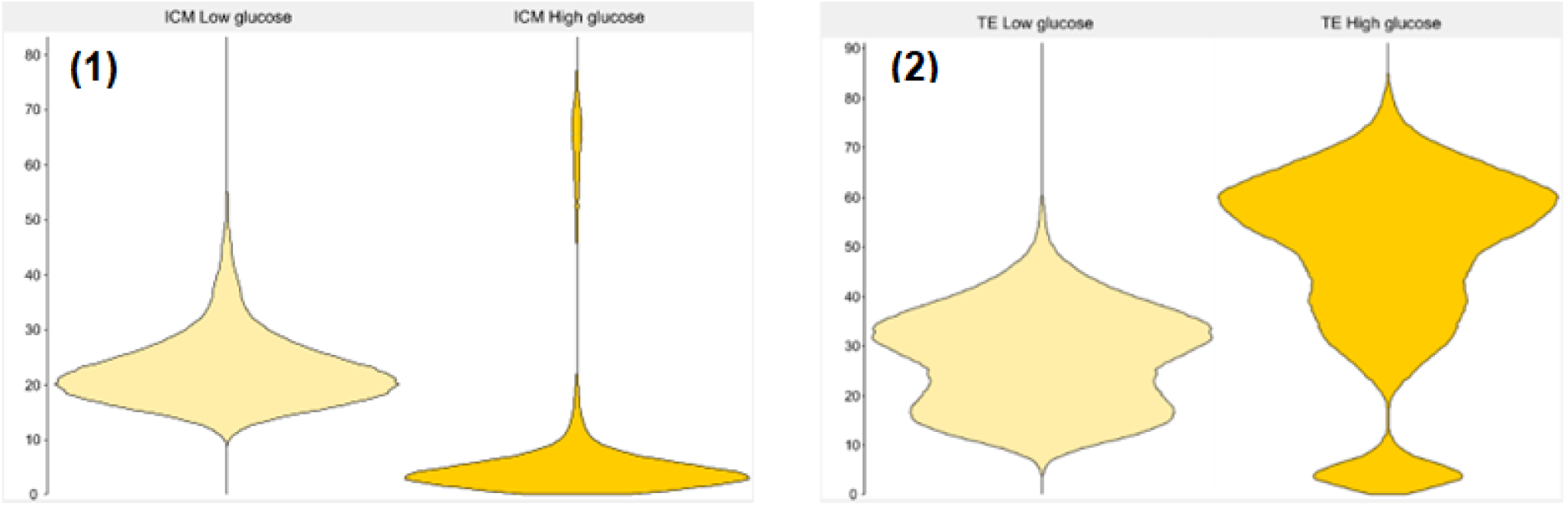
Mean Methylation levels in ICM and TE. (1): Bean plot showing DNA methylation average levels (0-100%) of grouped samples after EdgeR statistical test. Low glucose ICM (n=5) and high glucose ICM (n=5). (2): Bean plot showing TE DNA methylation average levels (0-100%) of grouped samples after EdgeR statistical test (<0.05). Low glucose TE (n=5) and high glucose TE (n=5).

5079 differentially methylated genes were identified; here, the most enriched gene sets were related to neurogenesis and morphogenesis. Regarding disease terms found using the OMIM database, cancer was most reported, followed by hypercholesterolemia, diabetes and Leigh syndrome. When using the GLAD4U database, the most enriched disease conditions were related to cell adhesion and mental disorders.

4265 hypermethylated genes were identified in the TE samples after culture in the high glucose condition. Here, the most enriched gene sets were neuron development, cell organisation and regulation of the plasma membrane. Regarding diseases identified from the OMIM database, cancer was most reported, followed by hypercholesterolemia, diabetes, hypertension and Leigh syndrome. When using information from the GLAD4U database, the most enriched diseases were again related to cell adhesion and mental disorders.

### High glucose ICM vs high glucose TE

The methylation analysis between ICM and TE samples in the high glucose condition resulted in 7 DMRs, which were overlapping 4 genes (UBE2D3, PIP5K1B, FAM122A and PLXNC1). Correlation analysis resulted in 0.945 and methylation levels were between 10-45%.

### Low glucose ICM vs low glucose TE

The methylation analysis between ICM and TE samples in the low glucose condition, resulted in 2 DMRs, which were overlapping 1 gene (DHX9). Correlation analysis resulted in 0.885 and methylation levels were between 10-45%.

### Differential methylation in H3K4me3 sites

Histone modifications are essential for regulating gene expression in development. Different types of histone modifications have diverse functions (Rivera and Ren, 2013); one example is trimethylated histone H3 lysine 4 (H3K4me3), which is known as an active histone modification (Sha et al., 2020). In the present study, it was found that DMRs from the high glucose group had considerably greater or lower levels of methylation (0-100%), whereas DMRs from low glucose samples were concentrated at mid-levels of methylation (data not shown).

Hypermethylated regions in the high glucose group (424), resulting in 314 gene IDs unambiguously mapped and 4 gene sets were enriched for the category of biological function. Moreover, genes CNOT3, CNOT6L (CCR4), DDX6, EDC4, EXOSC10 (RRP6), EXOSC3 (RRP40), LSM4, PFKL, XRN1 resulted as enriched for the RNA degradation pathway KEGG:03018, with an adjusted p-value of 0.007904.

### Differential methylation in H3K27me3 sites

Another histone modification, H3K27me3, is known to be a repressive histone mark, preferentially occupying the promoters of developmental genes (Xia *et al*., 2019). In the present study, potential repressor sites were mapped using the published report for ChIP H3K27me3 (accession number GSE124718). Called picks were used as a guide to select regions within the blastocyst genome that were further evaluated to obtain differentially methylated regions (DMRs).

Mean methylation levels from all H3K27me3 sites were analysed for low glucose and high glucose blastocysts. From these mapped regions, 4452 DMRs were identified in total, from which 588 were hypermethylated in low glucose and 3864 in high glucose.

On the other hand, hypermethylated regions in high glucose (3864) resulted in 1716 gene IDs that were unambiguously mapped for gene ontology analysis. This analysis resulted in 10 gene sets and 15 signalling pathways that were significantly (FDR) enriched; these were particularly related to neurological disorders, cardiovascular disorders and cancer.

### Differential methylation on CpG islands (CGI)

CpG islands were mapped for the homo sapiens genome GRCh38. Every CpG island was counted as a potentially methylated region, independent of the occurrence of CG or methylated points contained per island. The mean distribution of methylation levels in CpG islands has been represented in a trend plot (Fig. 7), displaying all mapped regions (22568 CpG islands) ±1Kb in low glucose and high glucose. In this study, we found that CpG islands were found to be hypermethylated in low glucose.

**Figure 7:**
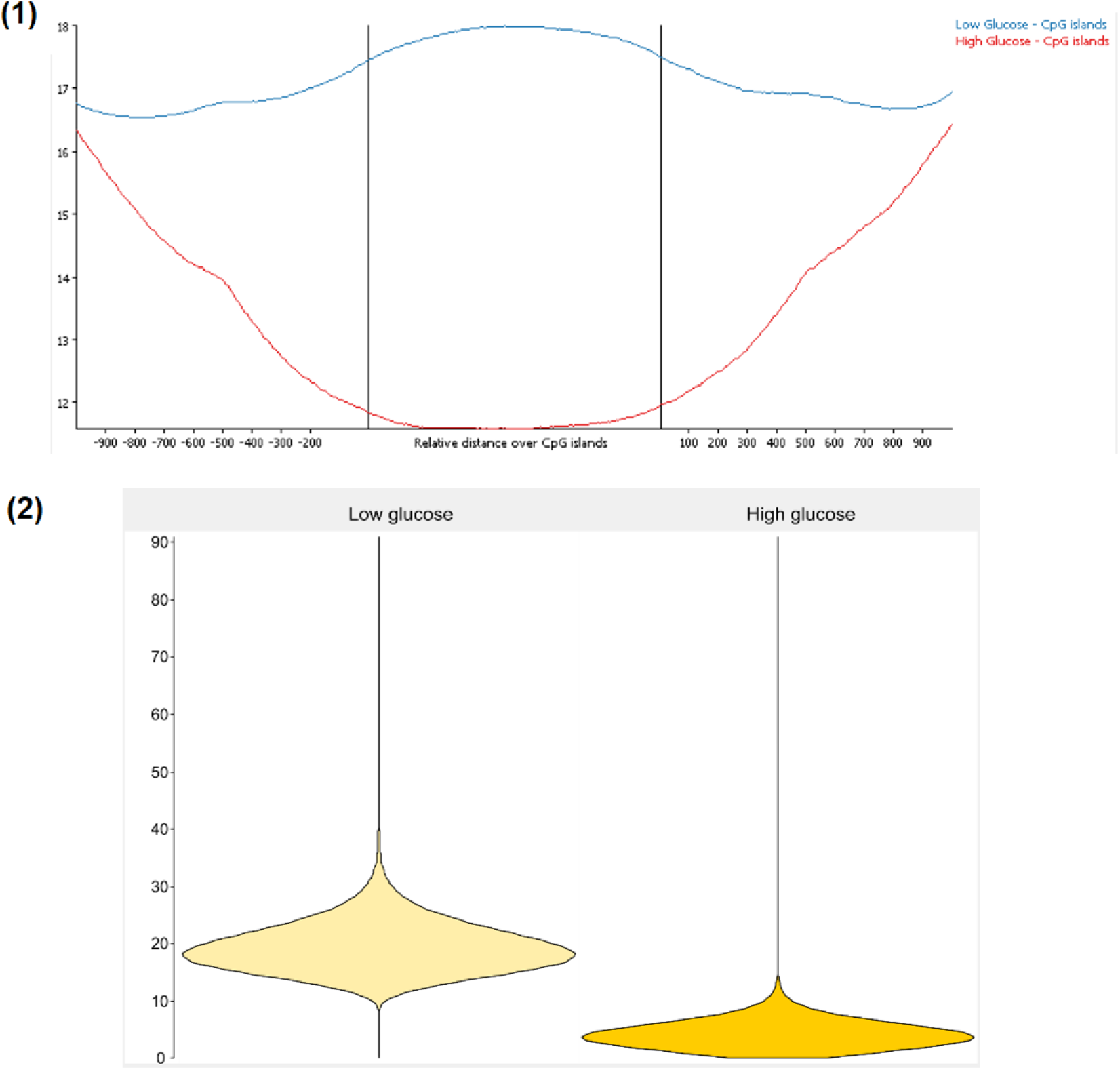
**(1):** Distribution of methylation levels in CpGi. Trend plot indicates the mean methylation levels from all mapped regions (22568 CpG islands) ±1Kb in low glucose (blue) and high glucose (red). **(2):** Methylation levels on CpG islands. Bean plots indicating percentage of methylation (Y-axis) and sample IDs (X-axis) for DMRs identified in this section, comparing distribution in high glucose vs low glucose. Differentially methylated regions (858) were identified between the two groups after applying Bismark quantification and EdgeR statistical analysis (p<0.05) after Benjamini and Hochberg correction, with a minimum difference of 20%.

**Figure 8.**
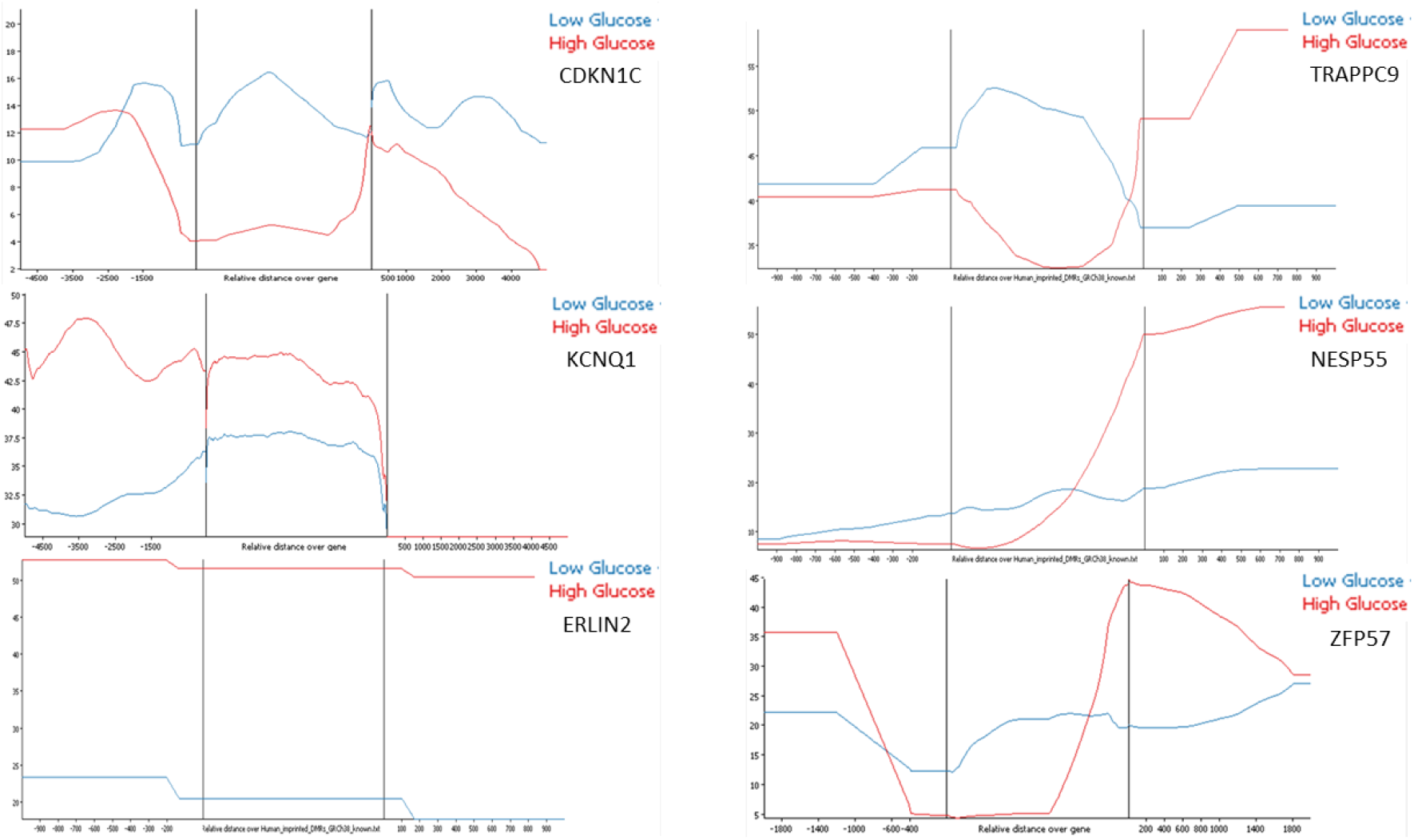
Quantitative trend plots of imprinted genes. Levels of methylation are distributed across the gene (upstream, gene body, downstream). Each gene showed different methylation levels (y-axis). CDKN1C from 2 to 20%, KCNQ1 from 30 to 50%, ERLIN2 from 20 to 50%, TRAPPC9 from 35 to 55%, NESP55 from 10 to 50%, ZFP57 from 5 to 45%. EdgeR for/rev Stats Filter on probes where High Glucose vs Low Glucose had a significance below 0.05 after Benjamini and Hochberg correction with a minimum difference of 10.0.

A total of 858 differentially methylated CpG islands (CPGi) were identified, which were found to overlap 780 genes. These have been identified with a significant increase in mean values of methylation for the low glucose blastocyst group, with a minimum difference of 20% relative to the high glucose group. Furthermore, examples of disease terms related to these genes included aplastic anaemia, pulmonary fibrosis, prostate cancer, mitochondrial complex I deficiency, leukaemia, acute myeloid leukemia, early infantile epileptic encephalopathy and autosomal dominant mental retardation, all based on OMIM and KEGG.

Hypermethylated regions in low glucose samples resulted in 560 genes, overlapped by CpGi sites. Hypermethylated regions in high glucose samples resulted in 220 genes, overlapped by CpGi sites. This resulted in the hypermethylated enrichment of the endocrine resistance pathway KEGG:01522, containing the following genes: ADCY6 (AC), CARM1 (CoA), GPER1 (GPR30), JAG2 (Jagged), NOTCH1 (Notch), SHC2 (Shc).

### Imprinted genes

Genomic imprinting is the parental allele-specific expression of certain genes (Morison et al., 2001). In normal development, imprinted genes play an essential role in regulating embryonic development and placental function. This process is typically regulated by differential DNA methylation inherited from the gametes (Varum *et al*., 2011).

The mean distribution of methylation in each imprinted gene was analysed, whereby mapped regions were represented as gene bodies ±1Kb. Trend plots allowed comparison of gene bodies, upstream and downstream regions, from low glucose and high glucose blastocysts (ICM+TE). CDKN1C demonstrated methylation levels between 2% and 20%, where the low glucose group was hypermethylated. KCNQ1 demonstrated methylation levels from 30% to 50%, where the high glucose group was hypermethylated. ERLIN2 showed methylation levels from 20% to 50%, where the high glucose group was hypermethylated. TRAPPC9 demonstrated methylation levels from 35% to 55%, where the low glucose group was hypermethylated (from the upstream region to gene body) and hypomethylated (in the downstream region). NESP55 presented methylation levels from 10% to 50%, where the low glucose group was hypermethylated (again from the upstream region to the gene body), and hypomethylated (in the downstream region). Finally, ZFP57 displayed methylation levels from 5% to 45%, where the high glucose group was hypermethylated in both the upstream and downstream region, and hypomethylated in the gene body.

## Discussion

ART has provided a unique insight into the molecular and metabolic processes that drive embryonic development, from a fertilised oocyte to an implantation-competent blastocyst. Previous research has compared different culture media, demonstrating that embryos can show differential gene expression and DNA methylation levels, when cultured in different commercially available culture media (Kleijkers *et al*., 2015).

As glucose is a standard component of human culture media, the present study was designed to determine whether changes in glucose concentration could influence global methylation and gene expression in the human preimplantation embryo. Using human embryos donated to research and cultured to blastocyst, we show that the concentration of glucose in human embryo culture medium is associated with changes in gene expression, global DNA methylation in both ICM and TE, and methylation levels at CpG islands and key histone modifications. These results are significant because glucose is a major nutrient metabolised by human embryos in culture, and yet we know relatively little of its downstream effects on the genome and epigenome. Since commercially available embryo culture media with varying glucose levels have also been associated with altered fetal growth, birthweight and postnatal development of IVF offspring, this work may have important ramifications for the long-term health of ART offspring.

By comparing global methylation of human blastocysts, we have been able to show that altering the glucose concentration in the culture media can cause significant changes to the global DNA methylome. In the blastocyst, there were 8654 DMGs between low and high glucose, with a high proportion (81.9%) of these being hypermethylated, with a smaller population (18.1%) being hypomethylated. In support of these findings, we also see that the DNA methyltransferase DNMT3A is upregulated in high glucose. DNA methyltransferases (DNMTs) are primarily responsible for placing methyl groups on CpG dinucleotides, leading to repression of gene expression. DNMT1 is primarily responsible for maintaining CpG methylation once these marks have been established. DNMT3A and DNMT3B, however, carry out de-novo DNA methylation (Canovas et al., 2017). In this study, DNMT3A was significantly upregulated in the high glucose condition. DNMT3A activity has previously been shown to play an important role in establishing DNA methylation during early embryo development and differentiation (Ho et al., 2016). Therefore, aberrations in the regulation of this gene could impact the epigenetic remodelling process. The TET enzymes (TET1, TET2, and TET3) oxidise 5mC to 5hmC, playing an epigenetic role in gene expression regulation (Matuleviciute et al., 2021). Together with passive loss of 5mC during DNA replication, TET enzymes play an essential role in the reprogramming of DNA methylation during early embryo development (Canovas *et al*., 2017). In the present study, the TET enzymes TET1 and TET2 were upregulated in the high glucose condition, relative to low glucose. Again, this indicates that DNA demethylation and remethylation in the preimplantation period is highly dynamic, and that glucose levels have the potential to impact early embryo gene regulation, potentially altering subsequent embryo viability (Canovas et al., 2017).

Whilst DNA methylation is generally repression of gene expression, we also note increased expression of key pluripotency genes such as Oct4 and Sox2 in high glucose, it is widely accepted that OCT4 is essential for maintaining pluripotency in the early mammalian embryo (Nichols et al., 1998). SOX2 and OCT4 also function together, balancing one another’s expression during embryo development (Rizzino and Wuebben, 2016). Regarding embryonic stem cell differentiation, the levels of OCT4 and SOX2 must be maintained within a narrow range, as small changes in these factors induce cellular differentiation (Rizzino, 2013). Thus, the significant downregulation of these two transcription factors in low glucose, could have downstream effects regarding the maintenance of pluripotency in the embryo, again potentially affecting embryo viability.

While the blastocyst as a whole shows 8654 DMGs, the ICM (2755 DMGs) and TE (5079 DMGs) are affected differently by glucose concentration. In the ICM. the great majority (2604) of DMGs were hypomethylated in high glucose, with only 151 being hypermethylated. In contrast, the TE DMGs resembled more the blastocyst DMGs, with 4265 genes hypermethylated and 814 hypomethylated, in high glucose.

The dissected TE cells appear to be more susceptible to epigenetic alterations, potentially affecting the normal development of the placenta. This would concur with reports in the literature, which state that changes in ART culture media have been associated with significant differences in birth weight of the offspring, and the placental weight to birth weight ratio, when compared with naturally conceived offspring (Eskild et al., 2013). The exact aetiology of how glucose could have this effect this is unclear, but as early embryo development is an essential period for the establishment and maintenance of epigenetic marks, any stress for the embryo during this period could result in perturbations in the epigenetic programming process, affecting gene expression throughout development.

### DNA methylation and potential for long term disease

In the present study, gene sets associated with later development of autism, mental disorders and epilepsy were found to be hypermethylated in embryos cultured in high glucose. Within the literature, aberrations in DNA methylation have been reported to play a role in the development of several neurological disorders (Xu and Li, 2012). The results reported from this study may indicate that exposure to glucose, above a certain level during early development, has the potential to alter the DNA methylation pattern of genes linked to neurological conditions. Although these findings seem concerning, it is important to state that the majority of published evidence thus far has confirmed equivalent neurodevelopment of ART-derived offspring, compared to naturally conceived offspring (Perros et al., 2022). However, longitudinal studies are still required to accurately confirm this for later life, so it is still vital to minimise any detrimental impacts from the early culture process.

The case is somewhat different regarding cardiometabolic measurements, where research has demonstrated differences in the cardiometabolic profile between IVF-conceived and naturally conceived offspring. Children born via ART appear to have higher systolic and diastolic blood pressures and higher fasting glucose levels, that could not be explained by alternate factors (Ceelen et al., 2009, Pontesilli et al., 2015). Epigenetic disturbances could be one mechanism contributing to this phenotype. To support this theory, the present study highlighted that gene sets associated with hypertension, hypercholesterolemia and diabetes were identified as hypermethylated in embryos cultured in high glucose. This supports the theory that alterations in the embryonic environment *in-vitro* could change metabolic gene expression during early development and that this could be a risk factor for later disease.

### Imprinted Genes

The present study has also demonstrated that methylation of imprinted genes can be altered by glucose concentration in embryo culture. Six imprinted genes (CDKN1C, KCNQ1, ERLIN2, TRAPPC9, NESP55, and ZFP57) were shown to be differentially methylated when comparing high and low glucose concentrations. Differentially methylated points were distributed through the gene body in a different manner for every gene and a clear pattern was not observed. Imprinting disorders are typically very rare in naturally conceived offspring, with incidence rates of around 1-10 per 100,000 births (Strawn et al., 2010). However, the incidence is higher in children born via ART, especially regarding Beckwith-Wiedemann syndrome (Henningsen et al., 2020); a growth disorder associated with pediatric cancer (Weksberg, 1994). Interestingly, alterations in the CDKN1C gene (Brioude et al., 2015) and loss of methylation of the imprinting centre region 2 within the KCNQ1 gene (Eßinger et al., 2020), have both been implicated in Beckwith-Wiedemann syndrome, and were both differentially methylated between glucose conditions in the present study. Moreover, different ontological analysis flagged the term ‘cancer’ or cancer related terms as significant in the hypermethylated and high glucose group. This provides evidence that genes involved in imprinting disorders can be affected by changes in glucose concentration, indicating a potential mechanism for the higher incidence of imprinting disorders seen in ART-derived offspring. Further studies are required to confirm if the changes seen in this study are maintained through cell divisions, during development and even trans-generationally.

In summary, ART has the potential to affect transcriptional regulation, metabolism and epigenetic variation. Minimal changes to the embryo culture environment can cause significant changes to metabolic pathways, which could lead to altered embryonic development. It is critical to continue research in this area, to work towards optimising culture conditions, minimising aberrant epigenetic alterations and minimising the risk profile to future offspring.

## References

Alpha Scientists in Reproductive Medicine and ESHRE Special Interest Group of Embryology (2011) ‘The Istanbul consensus workshop on embryo assessment: proceedings of an expert meeting’, Hum Reprod, 26(6), pp. 1270–83.

Arand, J., Reijo Pera, R. A. and Wossidlo, M. (2021) ‘Reprogramming of DNA methylation is linked to successful human preimplantation development’, Histochem Cell Biol, 156(3), pp. 197–207.

Brady, G., & Iscove, N. N. (1993). Construction of cDNA libraries from single cells. Methods in Enzymology, 225(1988), 611–623. 10.1016/0076-6879(93)25039-5

Brioude, F., Netchine, I., Praz, F., Le Jule, M., Calmel, C., Lacombe, D., Edery, P., Catala, M., Odent, S., Isidor, B., Lyonnet, S., Sigaudy, S., Leheup, B., Audebert-Bellanger, S., Burglen, L., Giuliano, F., Alessandri, J. L., Cormier-Daire, V., Laffargue, F., Blesson, S., Coupier, I., Lespinasse, J., Blanchet, P., Boute, O., Baumann, C., Polak, M., Doray, B., Verloes, A., Viot, G., Le Bouc, Y. and Rossignol, S. (2015) ‘Mutations of the Imprinted CDKN1C Gene as a Cause of the Overgrowth Beckwith-Wiedemann Syndrome: Clinical Spectrum and Functional Characterization’, Hum Mutat, 36(9), pp. 894–902.

Brison, D. R. and Leese, H. J. (1991) ‘Energy metabolism in late preimplantation rat embryos’, J Reprod Fertil, 93(1), pp. 245–51.

Cairo Consensus Group. 2020 ‘There is only one thing that is truly important in an IVF laboratory: everything’ Cairo Consensus Guidelines on IVF Culture Conditions. Reproductive BioMedicine Online. 40(1), pp.33–60. ISSN 1472-6483. 10.1016/j.rbmo.2019.10.003

Canovas, S., Ross, P. J., Kelsey, G. and Coy, P. (2017) ‘DNA Methylation in Embryo Development: Epigenetic Impact of ART (Assisted Reproductive Technologies)’, Bioessays, 39(11).

Ceelen, M., van Weissenbruch, M. M., Prein, J., Smit, J. J., Vermeiden, J. P., Spreeuwenberg, M., van Leeuwen, F. E. and Delemarre-van de Waal, H. A. (2009) ‘Growth during infancy and early childhood in relation to blood pressure and body fat measures at age 8-18 years of IVF children and spontaneously conceived controls born to subfertile parents’, Hum Reprod, 24(11), pp. 2788–95.

Dumoulin, J. C., Land, J. A., Van Montfoort, A. P., Nelissen, E. C., Coonen, E., Derhaag, J. G., Schreurs, I. L., Dunselman, G. A., Kester, A. D., Geraedts, J. P. and Evers, J. L. (2010) ‘Effect of in vitro culture of human embryos on birthweight of newborns’, Hum Reprod, 25(3), pp. 605–12

ESHRE (2021) Factsheet on infertility – prevalence, treatment and fertility decline in Europe. https://www.eshre.eu/-/media/sitecore-files/ESHREinternal/EU-Affairs/ESHRE_InfertilityFactsheet_v9.pdf

Eskild, A., Monkerud, L. and Tanbo, T. (2013) ‘Birthweight and placental weight; do changes in culture media used for IVF matter? Comparisons with spontaneous pregnancies in the corresponding time periods’, Hum Reprod, 28(12), pp. 3207–14.

Eßinger, C., Karch, S., Moog, U., Fekete, G., Lengyel, A., Pinti, E., Eggermann, T. and Begemann, M. (2020) ‘Frequency of KCNQ1 variants causing loss of methylation of Imprinting Centre 2 in Beckwith-Wiedemann syndrome’, Clin Epigenetics, 12(1), pp. 63.

Feuer, S. and Rinaudo, P. 2012. Preimplantation Stress and Development. Birth Defects Research Part C-Embryo Today-Reviews. 96(4), pp.299–314.

Fleming, T. P., Velazquez, M. A. and Eckert, J. J. (2015) ‘Embryos, DOHaD and David Barker’, J Dev Orig Health Dis, 6(5), pp. 377–83.

Gardner, D. K., Hamilton, R., McCallie, B., Schoolcraft, W. B. and Katz-Jaffe, M. G. (2013) ‘Human and mouse embryonic development, metabolism and gene expression are altered by an ammonium gradient in vitro’, Reproduction, 146(1), pp. 49–61.

Gardner, D. K. and Kelley, R. L. (2017) ‘Impact of the IVF laboratory environment on human preimplantation embryo phenotype’, J Dev Orig Health Dis, 8(4), pp. 418–435.

Ghosh, J., Coutifaris, C., Sapienza, C. and Mainigi, M. (2017) ‘Global DNA methylation levels are altered by modifiable clinical manipulations in assisted reproductive technologies’, Clin Epigenetics, 9, pp. 14.

Ho, S. M., Cheong, A., Adgent, M. A., Veevers, J., Suen, A. A., Tam, N. N. C., Leung, Y. K., Jefferson, W. N. and Williams, C. J. (2017) ‘Environmental factors, epigenetics, and developmental origin of reproductive disorders’, Reprod Toxicol, 68, pp. 85–104.

Håberg, S. E., Page, C. M., Lee, Y., Nustad, H. E., Magnus, M. C., Haftorn, K. L., Carlsen, E., Denault, W. R. P., Bohlin, J., Jugessur, A., Magnus, P., Gjessing, H. K. and Lyle, R. (2022) ‘DNA methylation in newborns conceived by assisted reproductive technology’, Nat Commun, 13(1), pp. 1896.

Henningsen, A.A, Gissler, M. Rasmussen, S. Opdahl, S. Wennerholm, U.B. Spangsmose, A.L. Tiitinen, A. Bergh, C. Romundstad, L.B. Laivuori, H. Forman, J.L. Pinborg, A. Lidegaard, Ø. (2020). Imprinting disorders in children born after ART: a Nordic study from the CoNARTaS group. Hum Reprod. 1;35(5):1178–1184. doi: 10.1093/humrep/deaa039.

Ivanova, E., Canovas, S., Garcia-Martínez, S., Romar, R., Lopes, J. S., Rizos, D., Sanchez-Calabuig, M. J., Krueger, F., Andrews, S., Perez-Sanz, F., Kelsey, G. and Coy, P. (2020) ‘Correction to: DNA methylation changes during preimplantation development reveal interspecies differences and reprogramming events at imprinted genes’, Clin Epigenetics, 12(1), pp. 96.

Kleijkers, S. H., Eijssen, L. M., Coonen, E., Derhaag, J. G., Mantikou, E., Jonker, M. J., Mastenbroek, S., Repping, S., Evers, J. L., Dumoulin, J. C. and van Montfoort, A. P. (2015) ‘Differences in gene expression profiles between human preimplantation embryos cultured in two different IVF culture media’, Hum Reprod, 30(10), pp. 2303–11.

Leese, H. J. (2012) ‘Metabolism of the preimplantation embryo: 40 years on’, Reproduction, 143(4), pp. 417–27.

Li, L., Lu, X. and Dean, J. (2013) ‘The maternal to zygotic transition in mammals’, Mol Aspects Med, 34(5), pp. 919–38.

Mandy, M. and Nyirenda, M. (2018) ‘Developmental Origins of Health and Disease: the relevance to developing nations’, Int Health, 10(2), pp. 66–70.

Matuleviciute, R., Cunha, P. P., Johnson, R. S. and Foskolou, I. P. (2021) ‘Oxygen regulation of TET enzymes’, FEBS J, 288(24), pp. 7143–7161.

Morbeck, D. E., Baumann, N. A. and Oglesbee, D. (2017) ‘Composition of single-step media used for human embryo culture’, Fertil Steril, 107(4), pp. 1055-1060.e1.

Morbeck, D. E., Krisher, R. L., Herrick, J. R., Baumann, N. A., Matern, D. and Moyer, T. (2014) ‘Composition of commercial media used for human embryo culture’, Fertil Steril, 102(3), pp. 759-766.e9.

Morison, I. M., Paton, C. J. and Cleverley, S. D. (2001) ‘The imprinted gene and parent-of-origin effect database’, Nucleic Acids Res, 29(1), pp. 275–6.

Nichols, J., Zevnik, B., Anastassiadis, K., Niwa, H., Klewe-Nebenius, D., Chambers, I., Schöler, H. and Smith, A. (1998) ‘Formation of pluripotent stem cells in the mammalian embryo depends on the POU transcription factor Oct4’, Cell, 95(3), pp. 379–91.

Perros, P., Psarris, A., Antsaklis, P., Theodora, M., Syndos, M., Koutras, A., Ntounis, T., Fasoulakis, Z., Rodolakis, A. and Daskalakis, G. (2022) ‘Neurodevelopmental Outcomes of Pregnancies Resulting from Assisted Reproduction: A Review of the Literature’, Children (Basel), 9(10).

Pinborg, A., Wennerholm, U. B., Romundstad, L. B., Loft, A., Aittomaki, K., Söderström-Anttila, V., Nygren, K. G., Hazekamp, J. and Bergh, C. (2013) ‘Why do singletons conceived after assisted reproduction technology have adverse perinatal outcome? Systematic review and meta-analysis’, Hum Reprod Update, 19(2), pp. 87–104.

Pontesilli, M., Painter, R. C., Grooten, I. J., van der Post, J. A., Mol, B. W., Vrijkotte, T. G., Repping, S. and Roseboom, T. J. (2015) ‘Subfertility and assisted reproduction techniques are associated with poorer cardiometabolic profiles in childhood’, Reprod Biomed Online, 30(3), pp. 258–67.

Rivera, C. M. and Ren, B. (2013) ‘Mapping human epigenomes’, Cell, 155(1), pp. 39–55.

Rizzino, A. (2013) ‘Concise review: The Sox2-Oct4 connection: critical players in a much larger interdependent network integrated at multiple levels’, Stem Cells, 31(6), pp. 1033–9.

Rizzino, A. and Wuebben, E. L. (2016) ‘Sox2/Oct4: A delicately balanced partnership in pluripotent stem cells and embryogenesis’, Biochim Biophys Acta, 1859(6), pp. 780–91.

Sha, Q. Q., Zhang, J. and Fan, H. Y. (2020) ‘Function and Regulation of Histone H3 Lysine-4 Methylation During Oocyte Meiosis and Maternal-to-Zygotic Transition’, Front Cell Dev Biol, 8, pp. 597498.

Smallwood, S. A., Lee, H. J., Angermueller, C., Krueger, F., Saadeh, H., Peat, J., Andrews, S. R., Stegle, O., Reik, W. and Kelsey, G. (2014) ‘Single-cell genome-wide bisulfite sequencing for assessing epigenetic heterogeneity’, Nat Methods, 11(8), pp. 817–820.

Strawn, E. Y., Bick, D. and Swanson, A. (2010) ‘Is it the patient or the IVF? Beckwith-Wiedemann syndrome in both spontaneous and assisted reproductive conceptions’, Fertil Steril, 94(2), pp. 754.e1-2.

Van Duijn, L., Steegers-Theunissen, R. P. M., Baart, E. B., Willemsen, S. P., Laven, J. S. E. and Rousian, M. (2022) ‘The impact of IVF culture medium on post-implantation embryonic growth and development with emphasis on sex specificity: the Rotterdam Periconceptional Cohort’, Reprod Biomed Online, 45(6), pp. 1085–1096.

Varum, S., Rodrigues, A. S., Moura, M. B., Momcilovic, O., Easley, C. A., Ramalho-Santos, J., Van Houten, B. and Schatten, G. (2011) ‘Energy metabolism in human pluripotent stem cells and their differentiated counterparts’, PLoS One, 6(6), pp. e20914.

Weksberg, R., Shuman, C. and Smith, A. C. (2005) ‘Beckwith-Wiedemann syndrome’, Am J Med Genet C Semin Med Genet, 137C(1), pp. 12–23.

Wilkinson, A. L., Zorzan, I. and Rugg-Gunn, P. J. (2023) ‘Epigenetic regulation of early human embryo development’, Cell Stem Cell, 30(12), pp. 1569–1584.

Wright, K., Brown, L., Brown, G., Casson, P. and Brown, S. (2011) ‘Microarray assessment of methylation in individual mouse blastocyst stage embryos shows that in vitro culture may have widespread genomic effects’, Hum Reprod, 26(9), pp. 2576–85.

Xia, W., Xu, J., Yu, G., Yao, G., Xu, K., Ma, X., Zhang, N., Liu, B., Li, T., Lin, Z., Chen, X., Li, L., Wang, Q., Shi, D., Shi, S., Zhang, Y., Song, W., Jin, H., Hu, L., Bu, Z., Wang, Y., Na, J., Xie, W. and Sun, Y. P. (2019) ‘Resetting histone modifications during human parental-to-zygotic transition’, Science, 365(6451), pp. 353–360.

Xu, Z., Li, X. (2012). ‘DNA Methylation in Neurodegenerative Disorders’ Curr Tran Geriatr Gerontol Rep 1, 199–205. 10.1007/s13670-012-0026-1

Zandstra, H., Brentjens, L. B. P. M., Spauwen, B., Touwslager, R. N. H., Bons, J. A. P., Mulder, A. L., Smits, L. J. M., van der Hoeven, M. A. H. B., van Golde, R. J. T., Evers, J. L. H., Dumoulin, J. C. M. and Van Montfoort, A. P. A. (2018) ‘Association of culture medium with growth, weight and cardiovascular development of IVF children at the age of 9 years’, Hum Reprod, 33(9), pp. 1645–1656.

